# *N*-linked glycan sites on the influenza NA head domain are required for efficient IAV incorporation and replication

**DOI:** 10.1101/2020.05.05.080077

**Authors:** Henrik Östbye, Jin Gao, Mira Rakic Martinez, Hao Wang, Jan-Willem de Gier, Robert Daniels

## Abstract

*N*-linked glycans commonly contribute to secretory protein folding, sorting and signaling. For enveloped viruses such as the influenza A virus (IAV), the addition of large *N*-linked glycans can also prevent access to epitopes on the surface antigens hemagglutinin (HA or H) and neuraminidase (NA or N). Sequence analysis showed that in the NA head domain of H1N1 IAVs three N-linked glycosylation sites are conserved and that a fourth site is conserved in H3N2 IAVs. Variable sites are almost exclusive to H1N1 IAVs of human origin, where the number of head glycosylation sites first increased and then decreased over time. In contrast, variable sites exist in H3N2 IAVs of human and swine origin, where the number of head glycosylation sites has mainly increased over time. Analysis of IAVs carrying N1 and N2 mutants demonstrated that the *N*-linked glycosylation sites on the NA head domain are required for efficient virion incorporation and replication in cells or eggs. It also revealed that N1 stability is more affected by the head domain glycans, suggesting N2 is more amenable to glycan additions. Together, these results indicate that in addition to antigenicity, N-linked glycosylation sites can alter NA enzymatic stability and the NA amount in virions.

## INTRODUCTION

Glycoproteins receive *N*-linked glycans when they are inserted into the endoplasmic reticulum (ER) lumen. Addition of these large oligosaccharide structures can lower the activation barrier for the glycoprotein to fold and can function as docking sites for cellular factors that assist in the folding, quality control and the trafficking of the newly synthesized glycoprotein [1–4]. Following the maturation process, *N*-linked glycans can also contribute to the function of the glycoprotein by influencing local conformations, extending the half-life, or directly participating in critical protein interactions [5–9]. Many envelope viral glycoproteins utilize *N*-linked glycans for these common cellular functions [10–12], and for the ability of the large glycan structures to limit access to sensitive epitopes [13–17]. For influenza viruses, the roles of *N*-linked glycans in the folding and masking of epitopes on its surface glycoprotein hemagglutinin (HA or H) have been well-established [13, 18–20]. However, a comprehensive picture is lacking for how *N*-linked glycans contribute to the other influenza glycoprotein neuraminidase (NA or N), as only a few experimental studies have been performed [17, 21].

The HA and NA glycoproteins from influenza A viruses (IAVs) are quite diverse and are classified into subtypes based on their antigenic and genetic properties [22]. Presently, sixteen HA (designated H1-H16) and nine NA (designated N1-N9) subtypes have been identified in avian IAVs in almost every possible combination [23]. Despite this variability, only H1N1 and H3N2 subtypes seasonally circulate in the human population, and they are also commonly isolated from swine species, which are susceptible to both avian and human IAVs [24, 25].

There are some similarities between the numerous NA subtypes. All are type II membrane glycoproteins that form a Ca^2+^-dependent tetrameric enzyme [26–30]. The enzymatic function, located in the C-terminal head domain, promotes the mobility of the virus by removing the terminal sialic acid residues that HA binds to on host and viral glycan structures [31, 32]. During IAV replication, NA is co-translationally targeted to the ER by its *N*-terminal transmembrane domain of varying hydrophobicity [27, 33, 34]. The transmembrane domain then inverts and integrates into the ER membrane as the C-terminal stalk and head domain are synthesized and translocated into the ER lumen [34]. Upon entering the Ca^2+^-rich ER lumen the stalk and the enzymatic head domain receive multiple *N*-linked glycans that are capable of recruiting chaperones [21, 35]. The chaperones likely assist in the folding and oligomerization of NA, which occurs through a cooperative process that involves the enzymatic head and the distal transmembrane domain [21, 36–38].

Previous work focused on the *N*-linked glycans of NA showed that an avian N9 variant predominantly misfolds in CHO cells when not glycosylated and that the misfolding is mainly caused by the loss of the head domain glycans [21]. More recent studies on H1N1 IAVs have begun to examine the temporal frequency of *N*-linked glycosylation sites in N1[39–41], and the heterogeneity of the *N*-linked glycan structures[35]. The positional analysis has led to the speculation that many N1 glycan sites correlate with antigenic regions[40], whereas the glycan analysis identified a single site on the N1 head domain that is modified by a wide variety of glycans, with a diverse antenna array when expressed in eggs[35]. However, the impact of these sites on NA has not been looked at directly for H1N1 IAVs and even less data is available for these sites from H3N2 IAVs.

Here, we examined the glycosylation site frequencies in the NA sequences from H1N1 and H3N2 strains by domain (stalk versus head), year of isolation and species of origin. Three conserved sites were identified in the N1 head domain and four in the N2 head domain. For the N1 head domain, variable glycosylation sites are almost exclusive to human H1N1 IAVs, whereas the N2 head domain has more variable sites and these are present in human, swine and avian H3N2 IAVs. Analysis of viruses carrying NAs with mutated glycosylation sites revealed that the ones in the head domain are required for efficient virion incorporation and replication, and influence NA stability in a subtype-dependent manner. These results illustrate how *N*-linked glycans on NA can perform multiple functions and may explain why variable glycosylation sites are more prevalent in N2.

## RESULTS

### Analysis of the N-linked glycan sites encoded by the NA from H1N1 IAVs

N-linked glycans are transferred to the Asn in the consensus sequence N-X-S/T-X, where X can be any amino acid other than Pro [42, 43]. The addition of the glycan occurs on the luminal side of the ER membrane (Fig. 1A), limiting the accessibility of some regions in membrane proteins. Influenza NA is synthesized as a type II membrane glycoprotein with the N-terminus in the cytosol and a long C-terminal region in the ER lumen (Fig. 1B). The C-terminal region contains the stalk and head domain, and both of these regions have been shown to encode multiple *N*-linked glycosylation sites [21, 34, 39–41]. To determine if the glycosylation sites in NA have a domain or temporal bias with respect to the IAV species of origin, we initially examined the available subtype 1 (N1) sequences from H1N1 IAVs. The sequence analysis showed that the majority (∼92%) of avian N1s possess seven predicted *N*-linked glycan sites, whereas the human and swine N1s tend to have more sites and vary in the site number (Fig. 1C, left panel). In the stalk, most avian N1s have four sites and the human and swine N1s mainly carry four or five (Fig. 1C, middle panel). In the head domain, avian N1s predominantly have three sites, whereas the swine N1s contain either three or four, and the human N1s range from three to five (Fig. 1C, right panel). In line with previous reports [39, 41], these differences indicate that the stalk and head domain both contribute to the species-related variation in the number of NA glycosylation sites, but it remains unclear if the bias relates to how the sequences were collected.

**Figure 1.**
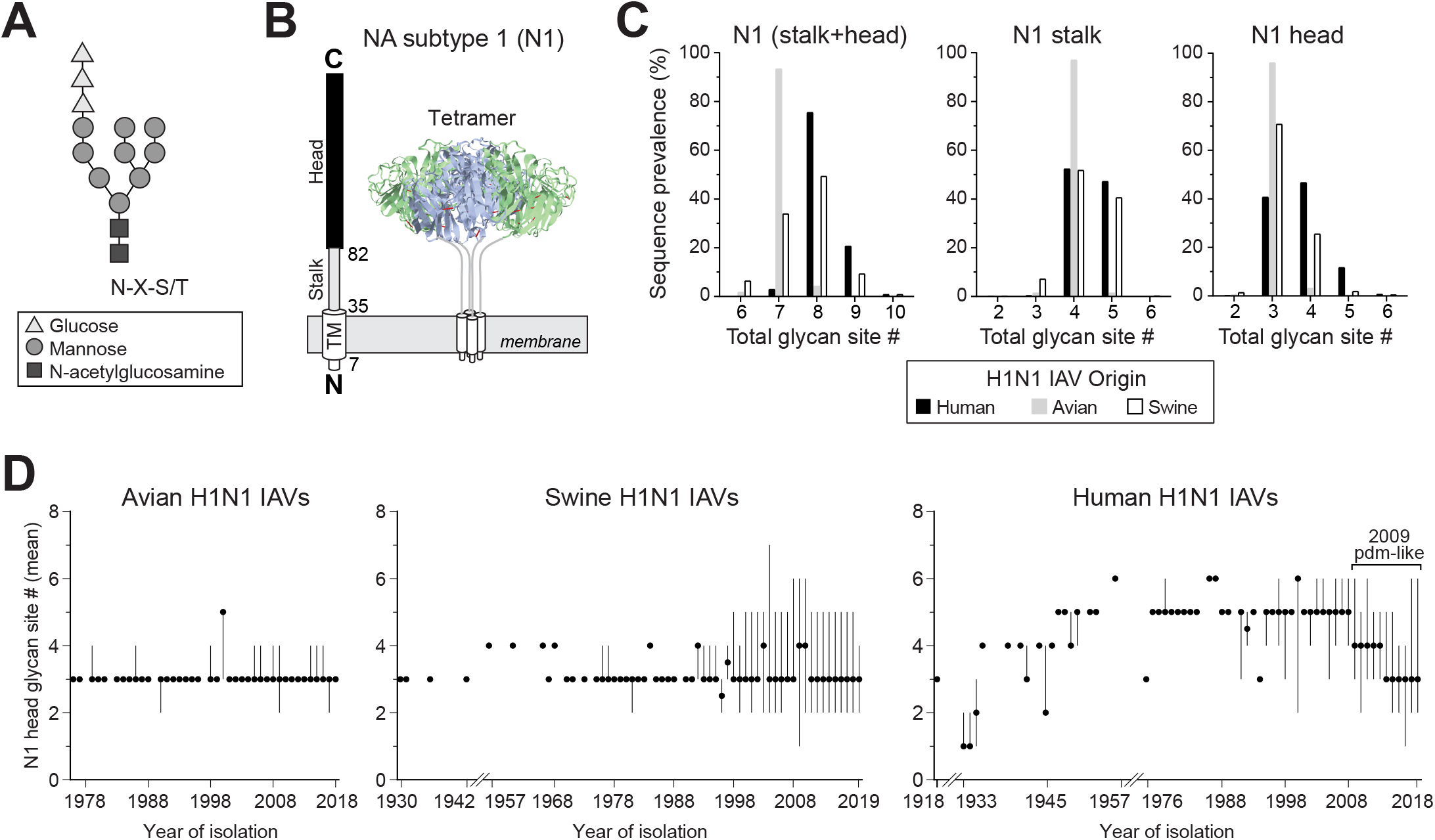
*N*-linked glycan site variation in NA from H1N1 IAVs by species of origin and time. **A.** Diagram of an *N*-linked glycan structure that is transferred to secretory glycoproteins during entry into the ER lumen. The glycan is added to the Asn (N) of the consensus sequence N-X-S/T. **B.** Linear and structural organization of the domains in NA from H1N1 IAVs. The numbers correspond to the amino acid position at the start of the transmembrane (TM), stalk and head domains of these NAs. **C.** Graphs showing the prevalence of the N1 sequences from human (n=18966), swine (n=4949), and avian (n=630) H1N1 IAVs that possess the indicated number of glycosylation sites in the stalk and head domain (**left panel**), the stalk alone (**middle panel**) and the head domain alone (**right panel**). **D.** Temporal graphs displaying the mean number of glycosylation sites in the N1 head domain with respect to the year the avian (**left panel**), swine (**middle panel**), and human (**right panel**) H1N1 IAVs were isolated. Filled circles correspond to the mean. Lines show the range in the number of sites in the sequence set for each year. All analyses were performed using full-length NA sequences downloaded from the NCBI Influenza Database.

Some glycosylation sites in the NA head have been linked to antigenicity [17], therefore we performed a temporal analysis of the sites in the N1 head domain. Regardless of the year the strain was isolated, the avian N1 head domains were found to mainly contain three sites (Fig. 1D, left panel) and the swine N1 head domains fluctuated between three to four sites with no temporal pattern (Fig. 1D, middle panel). In contrast, the human N1 head domains showed a step-wise pattern, with the early strains increasing in the number of sites from three to six, and the more recent strains decreasing back to three (Fig. 1D, right panel). These temporal observations indicate that the addition and removal of *N*-linked glycan sites in the N1 head domain is more characteristic of human H1N1 IAVs and that N1 likely requires at least three glycosylation sites in the head domain.

### Location of the N-linked glycan sites in the NA head domain of H1N1 IAVs

Positional analysis revealed that three *N*-linked glycosylation sites are highly conserved in the N1 head domain in H1N1 IAVs (Fig. 2A). *In silico* modelling of minimal glycan structures onto these sites showed that one (Asn146) is positioned on the top of the NA tetramer and the other two (Asn88 and Asn235) are located close together on the bottom, facing the viral membrane (Fig. 2B). One of the four prevalent variable sites in the human N1 head domain (Asn386) is also frequently found in the swine N1 head domain (Fig. 2C), likely due to the swine origin of the human 2009 pandemic H1N1 virus. Temporally, the prevalent variable sites overlap for different time periods, contributing to the discrete changes observed in the number of glycosylation sites on the human N1 head domain (Fig. 2C and 1F). Positionally, three of the variable sites (Asn365, Asn386, and Asn455) cluster towards the NA tetramer side (Fig. 2D), which has previously been shown to be an antigenic region in NA [44]. The final prevalent variable site at Asn434 is located very close to the conserved site at Asn146 (Fig. 2D), suggesting these two sites, and possibly the other two conserved sites (Asn88 and Asn235), perform redundant roles.

**Figure 2.**
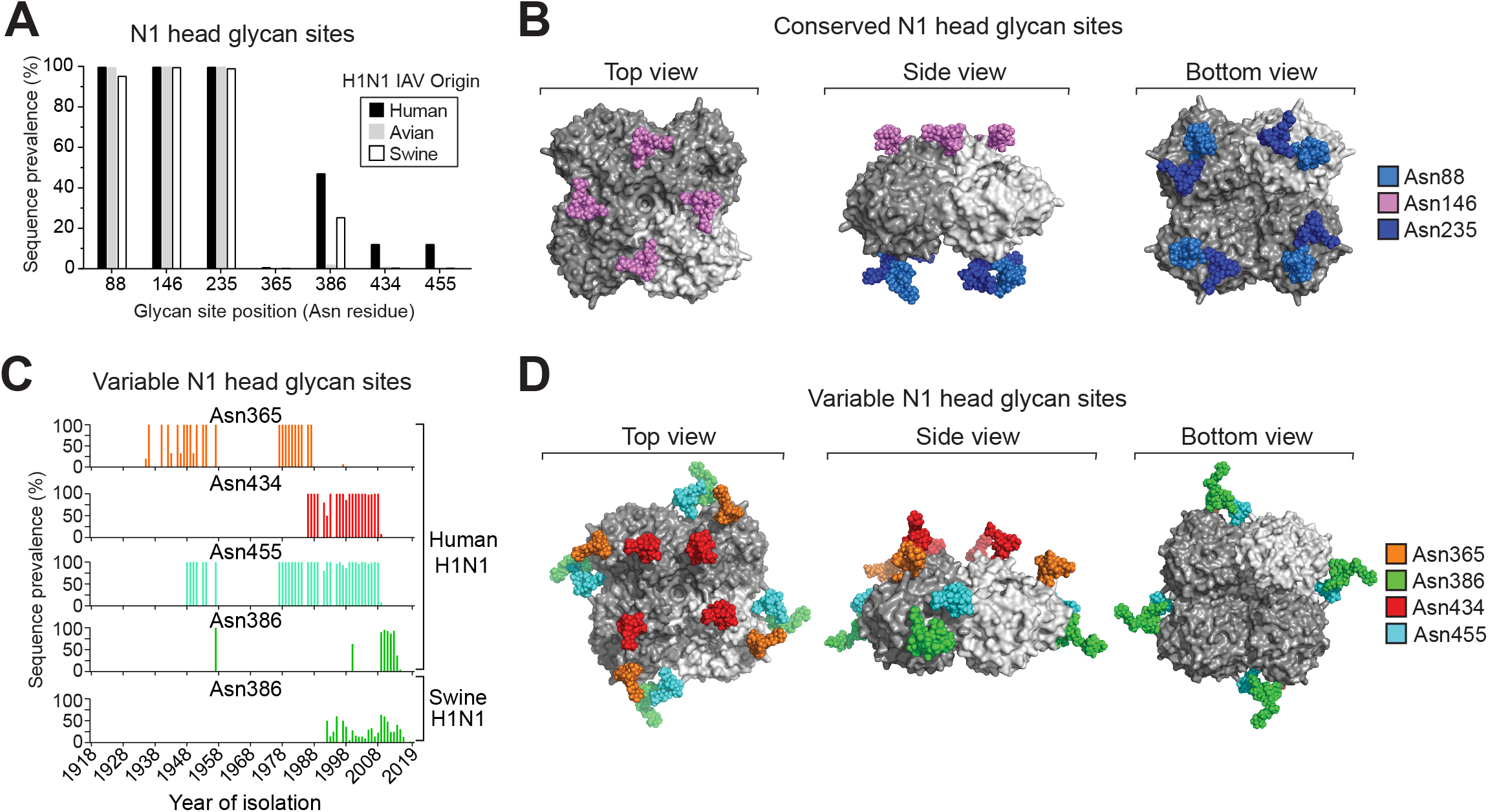
Positions of the *N*-linked glycan sites in the NA head domain from H1N1 IAVs. **A.** Graph showing the prevalence of the most frequent head domain glycosylation sites in the N1 sequences based on the H1N1 IAV species of origin. The positioning refers to the Asn (N) in the N-X-S/T sequence. **B.***In silico* model of *N*-linked glycan structures mapped onto the conserved sites of a 2009 pandemic-like N1 head domain structure (PDBID: 5NWE) [53]. **C.** Temporal graphs displaying the frequency of the most prevalent variable glycosylation sites in the head domain of N1 with respect to the year the sequences were isolated. **D.***In silico* model of *N*-linked glycans mapped onto a N1 head domain structure (PDBID: 5NWE) [53] at amino acids that correspond to the position of the prevalent variable head glycosylation sites.

### The conserved N1 head glycosylation sites are not essential for viral replication in cells

Although the glycosylation sites at Asn88, Asn146 and Asn235 are highly conserved in the N1 head domain (Fig. 3A), we were able to identify sequences that carry a mutation in one of the sites (S90P, T148A, and N235K). These natural mutations were then introduced into a NA (N1-MI15) from a previously recommended 2009 pandemic-like H1N1 vaccine strain (A/Michigan/45/2015) in various combinations to determine if the conserved sites are required for H1N1 IAV replication. We chose N1-MI15 for the analysis as it does not contain any variable head glycan sites. An additional mutation (S90A) was also included to alleviate potential folding concerns associated with the S90P mutation that introduces a Pro residue.

**Figure 3.**
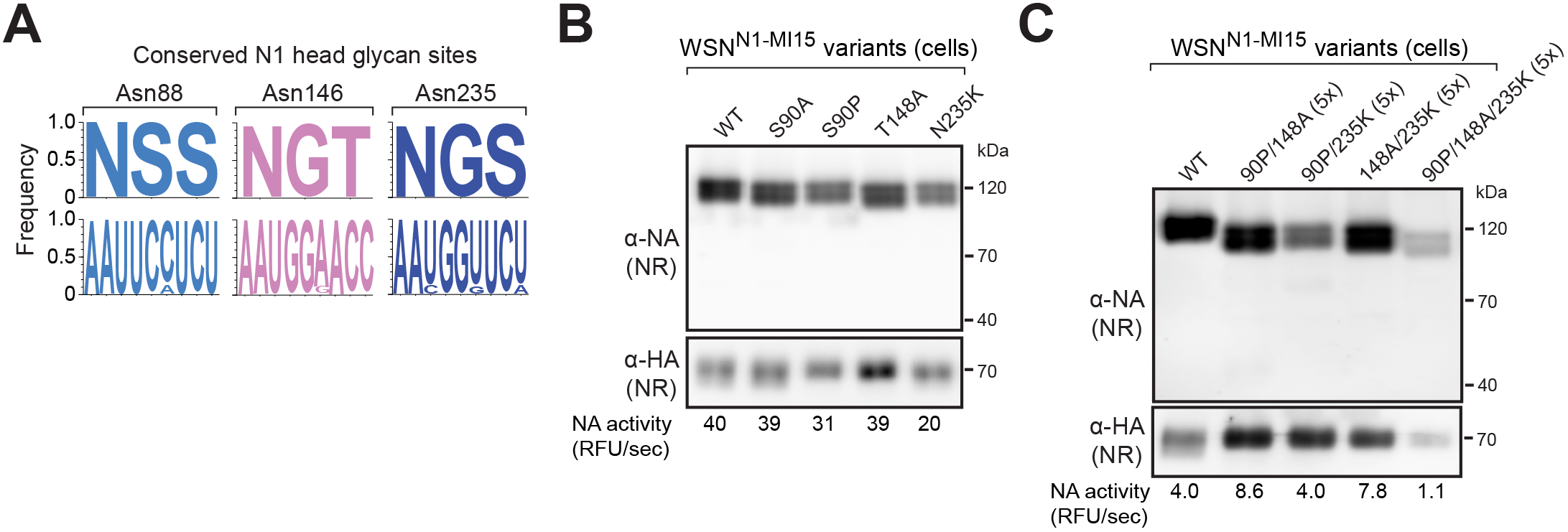
The conserved N1 head glycosylation sites are not essential for viral replication in cells. **A.** Logo plots displaying the amino acid (top) and nucleotide (bottom) frequency for the three conserved glycosylation sites on the N1 head domain from human H1N1 IAVs. **B and C.** Representative NA and HA immunoblots of recombinant WSN viruses carrying N1-MI15 with the indicated glycosylation site mutations. The viruses were rescued by reverse genetic, passaged in MDCK cells and the viral-containing supernatants were sedimented and resuspended in equal volumes. Each sample was split, one part was resolved by non-reducing (NR) SDS-PAGE prior to immunoblotting, and the other was used to determine the NA activity values, which are listed below the blots. Samples (**C**) containing resuspension volumes greater than the WT control are indicated as a ratio in the parenthesis.

Surprisingly, all the recombinant viruses carrying the N1-MI15 single, double and triple glycan site mutants were rescued using a WSN33 backbone. These single-gene reassortant viruses, along with a N1-MI15 wild-type (WT) control, were propagated in MDCK cells, isolated by sedimentation, and analysed. Each of the N1-MI15 variants in the sedimented virions possessed enzymatic activity and resolved as intermolecular disulphide bonded dimers following non-reducing (NR) SDS-PAGE (Figs. 3B and 3C), which is a characteristic of properly folded N1 [26, 36]. The expected mobility increase was more pronounced for N1-MI15 with the triple and double glycosylation site mutants than for the single site mutants (Figs. 3B and 3C), implying the appropriate glycans were likely absent in the mutants. We also noted that visualization of the double and triple glycan site mutants required higher volumes of the sedimented viral-containing medium (Fig. 3C), suggesting the absence of several conserved glycans modestly impairs viral growth by decreases NA folding or trafficking.

### The conserved glycosylation sites in the N1 head are not essential for viral replication in eggs

Several factors can potentially influence the results obtained after the conserved glycan sites were mutated in N1-MI15, including the use of the natural mutations, the particular NA, the viral backbone and the growth environment. Therefore, we repeated the analysis with a NA (N1-BR18) from a more recently recommended 2009 pandemic-like H1N1 vaccine strain (A/Brisbane/02/2018) using a different mutation strategy (N to Q), backbone (PR8) and growth environment (embryonated eggs). Like the prior results, all the recombinant viruses carrying N1-BR18 with single, double and triple mutations in the head glycan sites were rescued. Upon passaging in eggs lower hemagglutinating unit (HAU) titres were only observed for the two double glycan site mutants lacking the Asn146 site and the triple glycan site mutant (Fig. 4A, upper graph). Variable NA activity was measured in the egg allantoic fluid for all the viruses apart from the triple glycan site mutant, which consistently produced low activity levels (Fig. 4A, lower graph). Each of the N1-BR18 mutants displayed increased mobility on reducing (RD) and NR SDS-PAGE, which correlated with the number of the glycosylation site mutations (Fig. 4B), indicating the mutations remained intact. Together, these results demonstrate that the conserved *N*-linked glycosylation sites on the N1 head domain are not essential for H1N1 virus replication in cells or embryonated eggs. However, we did observe that viral replication appears to decrease when the three conserved head glycan sites are mutated (Fig. 4A) and that N1 intermolecular disulphide-bond formation and virion incorporation are less efficient when two glycan sites are absent (Fig. 4B).

**Figure 4.**
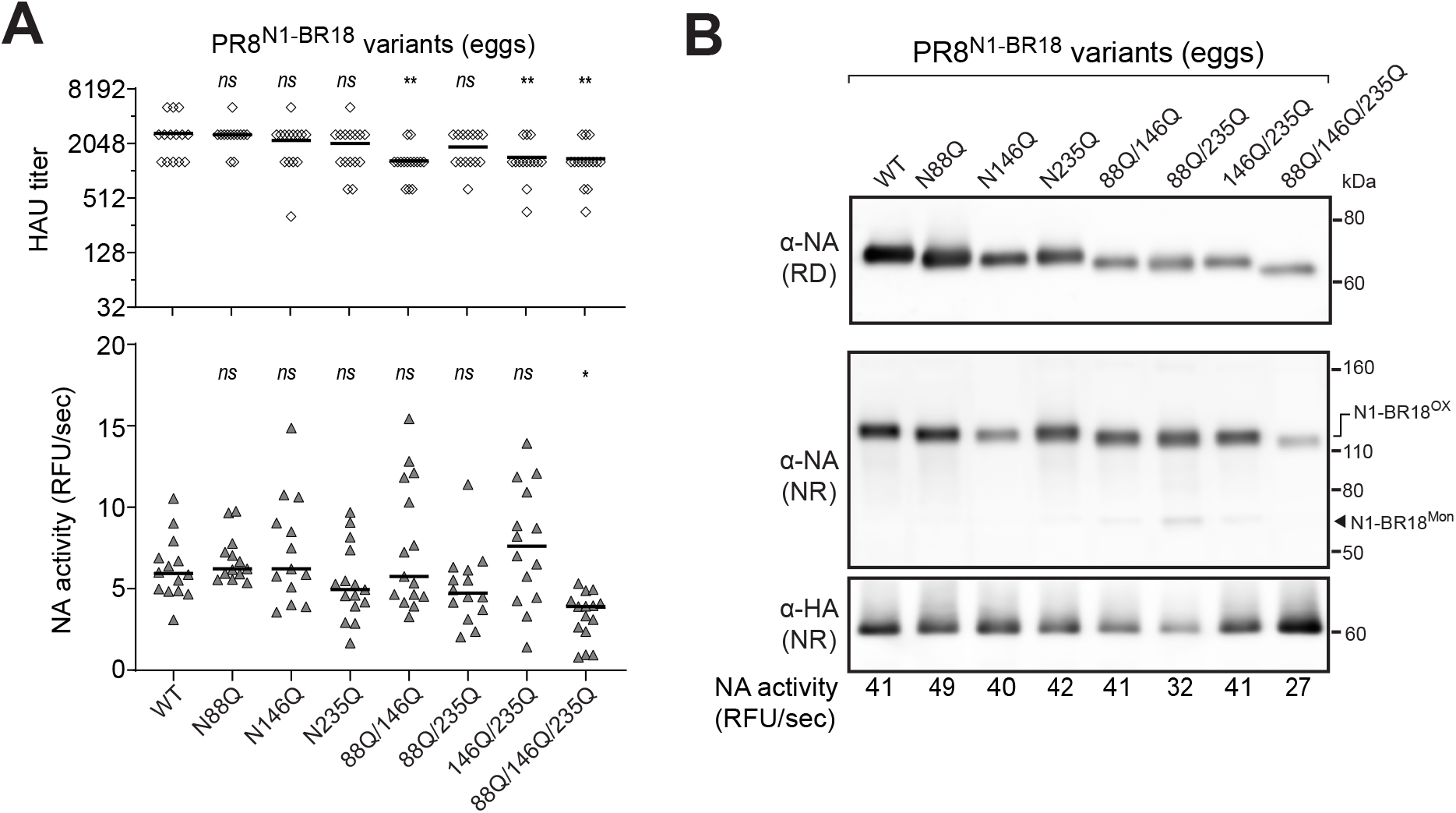
Mutation of the conserved N1 head glycosylation sites causes slight viral replication defects in eggs. **A.** Scatter plots of the HAU titers and NA activities that were measured in the allantoic fluid harvested from eggs infected with recombinant PR8 viruses carrying N1-BR18 with the indicated glycosylation site mutations. The viruses were rescued by reverse genetics and passaged twice in eggs. The data points from individual eggs following the second passage are shown together with the median (line). *P* values (95% CI) were determined with respect to the WT values by a one-way ANOVA. **B.** Representative NA and HA immunoblots of the recombinant PR8^N1-BR18^ viruses with the indicated mutations in the N1 head glycosylation sites. The allantoic fluid from the second passage was pooled, the virions were isolated by centrifugation and adjusted to equal total protein concentration prior to being resolved by NR and reducing (RD) SDS-PAGE. NA activity (below the immunoblots) in the virions was also measured using equal total protein amounts.

### The conserved glycans on the head domain influence N1 viral incorporation

When the three conserved glycan sites in the N1-BR18 head domain were mutated, the PR8 backbone virus showed a tendency in eggs to reach lower HAU titres and NA activity levels (Fig. 4A). This raised several questions: is the phenotype backbone dependent; do the mutations change the HA to NA ratio in the virions, or cause a decrease in viral production? To address these questions, we rescued a wildtype N1-BR18 virus (WT) and a mutant containing no head glycosylation sites (NHG 3Q) using a WSN backbone. Upon passaging in eggs both the HAU titre and the NA activity in the allantoic fluid were significantly lower when the three conserved glycosylation sites on the N1 head domain were mutated (Fig. 5A, compare WT to NHG 3Q), demonstrating the phenotype is conserved and may be somewhat exacerbated with a WSN backbone.

**Figure 5.**
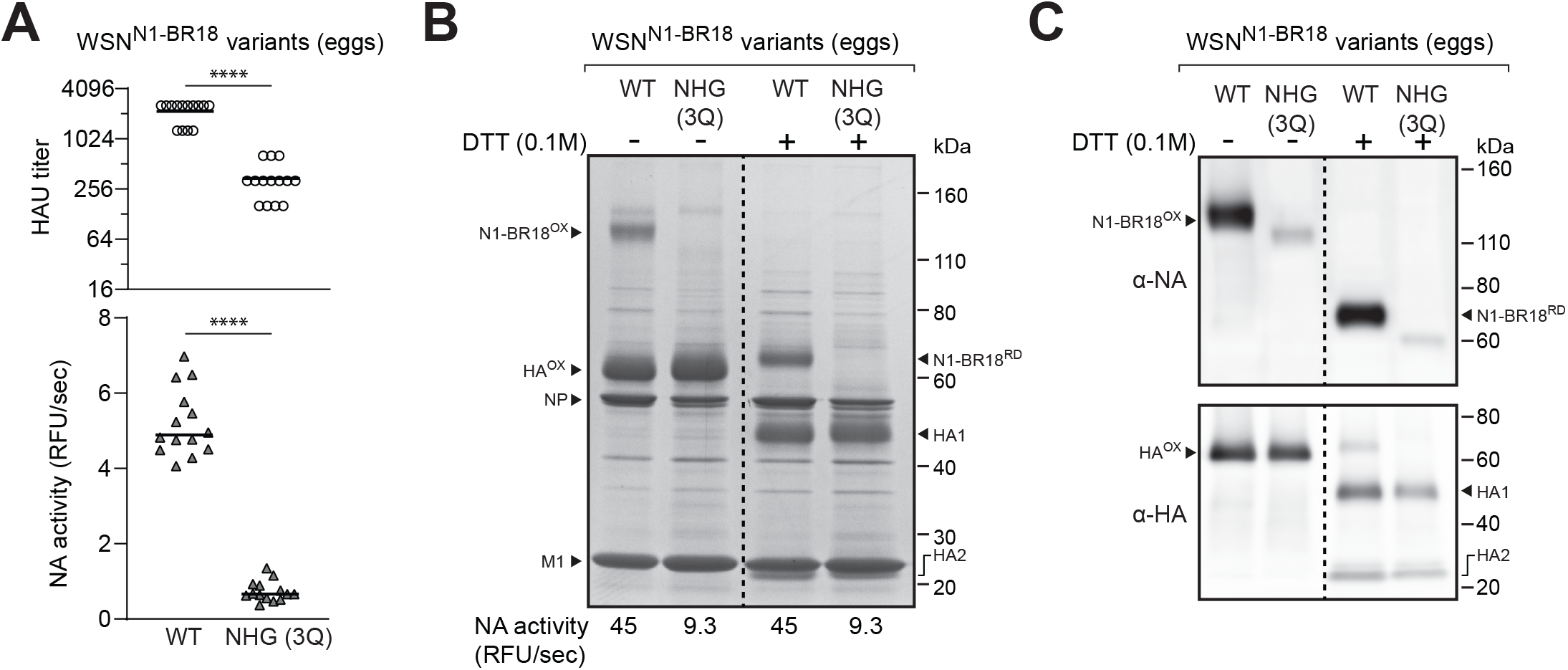
N1 viral incorporation is reduced when the conserved head glycosylation sites are absent. **A.** Scatter plots of the HAU titers and NA activities in the allantoic fluid harvested from eggs infected with recombinant WSN viruses carrying N1-BR18 (WT) or N1-BR18 with no head glycan sites (NHG 3Q), which was generated by Gln mutations of each head glycosylation site (N88Q, N146Q and N235Q). The viruses were rescued by reverse genetic s and passaged twice in eggs. The data points from individual eggs following the second passage are shown with the median. *P* values (95% CI) are from a two-tailed unpaired t-test. **B.** Representative image of a Coomassie stained SDS-PAGE gel containing the indicated recombinant WSN^N1-BR18^ viruses. The allantoic fluid from a second passage in eggs was pooled, the virions were isolated by centrifugation, and the protein concentration was determined. Samples containing ∼5μg of total protein were treated with the reductant DTT as indicated and resolved using a 4-12% Tris-glycine SDS-PAGE gel. Inter- (N1-BR18^OX^) and intra- (HA^OX^) molecular disulfide bonded NA and HA are indicated along with the reduced forms (N1-BR18^RD^, HA1, and HA2). The viral proteins NP and M1 are also indicated. The NA activity listed below the gel was measured using equal total protein amounts of the two viruses. **C.** NA and HA immunoblots of the isolated recombinant WSN^N1-BR18^ viruses. Samples containing equal total protein amounts were treated with DTT as indicated, resolved using a 4-12% Tris-glycine SDS-PAGE gel and transferred to a PVDF membrane prior to immunoblotting.

The lower HAU titres for the virus carrying the N1-BR18 mutant (NHG 3Q) indicated that the conserved glycosylation sites on the N1 head domain do contribute to viral production. To address if the mutations changed the HA to NA ratio in the virus, we isolated the virions by centrifugation and examined equal quantities of total protein by SDS-PAGE followed by Coomassie staining. In the absence of the reductant dithiothreitol (DTT), oxidized N1-BR18 dimers were readily apparent for the WT virus and these were reduced to the expected molecular weight upon DTT addition (Fig. 5B). Despite the relatively equivalent levels of HA, NP and M1, a band corresponding to N1-BR18 with the three glycan site mutations (NHG 3Q) was not observed, indicating the HA to NA ratio increased (Fig. 5B). Although the N1-BR18 mutant was not visible by Coomassie staining, the virus displayed NA activity levels ∼20% of the WT when equal viral protein amounts were analysed (Fig. 5B) and the protein was also detected as a less intense faster migrating band by immunoblotting (Fig. 5C). Together, these results demonstrate that H1N1 viral production and NA incorporation both decrease when all three conserved glycosylation sites are absent on the N1 head domain.

### The variable glycosylation sites in the N1 head domain from human H1N1 IAVs

Previous studies have demonstrated that the *N*-linked glycosylation sequence N-X-T is more efficiently recognized than N-X-S [43]. Based on sequence alignments of the human N1 head domain three of the main variable glycosylation sites (Asn365, Asn386 and Asn455) use N-X-S and these sites likely change by substitutions at either the N or S residues (Fig. 6A). In contrast, the other variable glycosylation site (Asn434) uses N-X-T and appears to have been created by a T codon insertion combined with a N substitution that occurred later (Fig. 6B). The codon insertion is almost exclusively found in human H1N1 IAVs beginning in 1948 and ending when the 2009 pandemic H1N1 virus, which carries a N1 gene segment of swine origin, was introduced to the human population (Fig. 6B). Positionally, the Asn434 site is located near the conserved Asn146 site and out of the seven most prevalent glycosylation sites, these are the only two with the more efficient recognition sequence, suggesting they may impact N1 more than the others.

**Figure 6.**
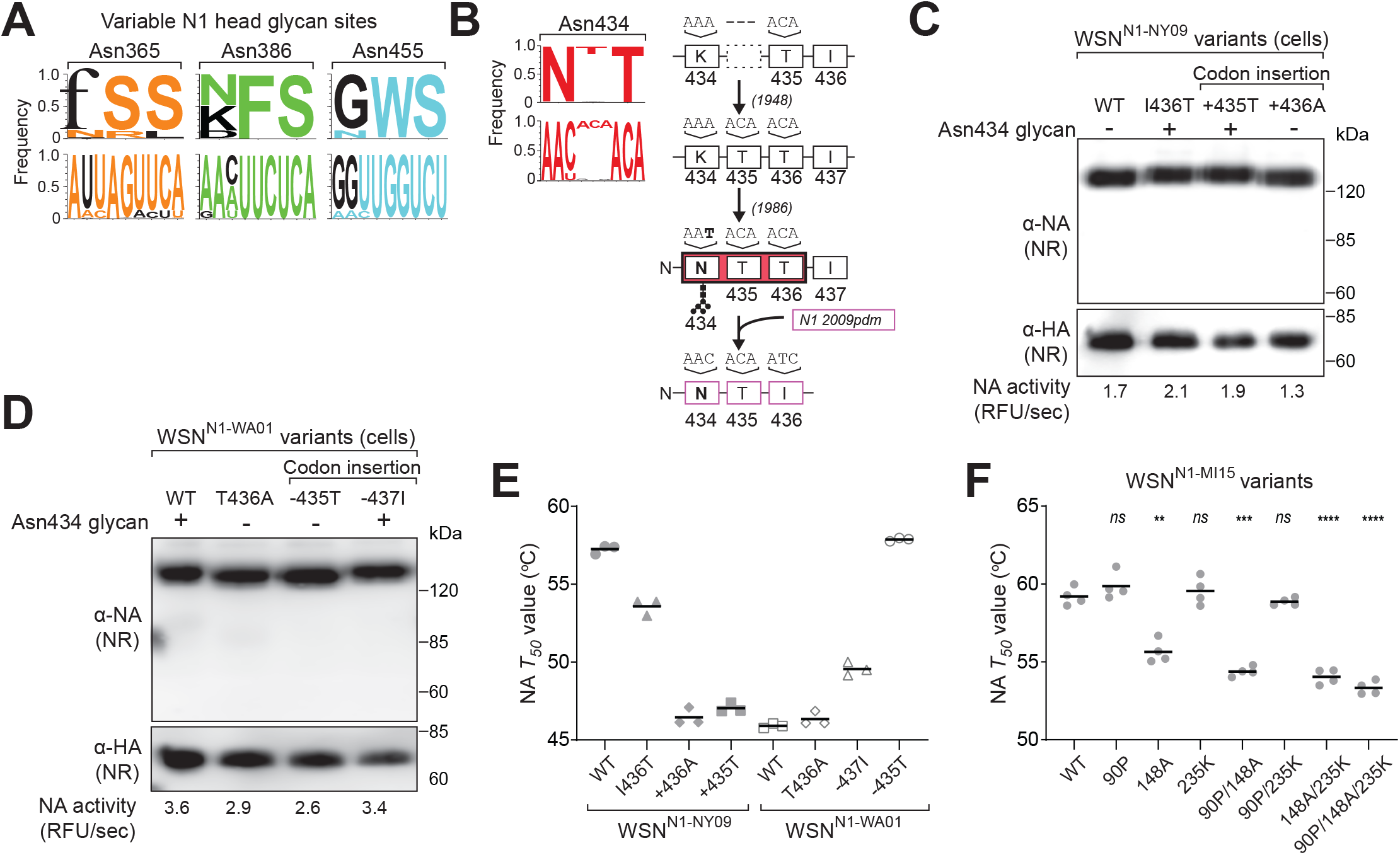
N1 stability decreases from the head domain insertion that creates the variable Asn434 glycosylation site. **A.** Logo plots displaying the amino acid (top) and nucleotide (bottom) frequency for the three N-X-S variable glycosylation sites on the N1 head domain from human H1N1 IAVs. **B.** Amino acid (top) and nucleotide (bottom) logo plot for the N-X-T variable glycosylation site in the N1 head domain from human H1N1 IAVs. A timeline showing when the insertion and the necessary mutations appeared in the database is included to the right. Note that the N1 ‘insertion’ was lost in 2009 when the pandemic H1N1 IAV of swine origin became prevalent. **C and D.** Representative NA and HA immunoblots of recombinant WSN viruses carrying (**C**) N1-NY09 with the indicated mutations and insertions or (**D**) N1-WA01 with the indicated mutations and deletions. The viruses were rescued by reverse genetic, passaged in MDCK cells and the viral-containing supernatants were sedimented, resuspended in equal volumes and resolved by NR SDS-PAGE. **E.** NA *T*_50_ temperatures are displayed for the recombinant WSN viruses carrying N1-NY09 or N1-WA01 with the indicated mutations, insertions or deletions. The measurements (n = 3 biologica lly independent experiments) were taken in tissue culture medium with the NA activity at 37°C set to 100%. The line represents the mean. **F.** NA *T*_50_ temperatures are displayed for the WSN reassortant viruses carrying N1-MI15 with the indicated glycan site mutations. The measurements (n = 4 biologically independent experiments) were taken in tissue culture medium with the NA activity at 37°C set to 100%. The line depicts the mean and the *P* values (95% CI) were determined with respect to WT by a one-way ANOVA.

To investigate this site, two similar natural sequences were identified that lack or have the insertion resulting in the Asn434 glycosylation site. Several mutations were then introduced into the NA lacking the insertion (N1-NY09), which is from a 2009 pandemic-like H1N1 strain (A/New York/18/2009), and a NA possessing the insertion (N1-WA01), which is from a 2001 seasonal H1N1 strain (A/Waikato/7/2001). For N1-NY09 these involved creating a glycosylation site by mutation (I436T), codon insertion (+435T), and a control where a codon insertion (+436A) was made that does not create a glycosylation site. All the mutants were rescued using a WSN backbone and propagated using MDCK cells where no growth defect was observed based on cytopathic effects. Activity measurements and immunoblot analysis of the isolated particles showed no significant change in the N1-NY09 levels and the mutants with the additional glycosylation site (I436T and +435T) displayed the expected mobility increase (Fig. 6C). Similar results were obtained from viruses carrying N1-WA01 with converse mutations that removed the glycosylation site (T436A), the codon insertion responsible for the glycosylation site (−435T) or a downstream codon (−437I) that left the site (Fig. 6D).

As Asn434 is near the central Ca^2+^ binding site, which is a major determinant for NA stability [26], we asked if this glycan influences NA thermostability. For N1-NY09, introducing the insertion and the glycosylation site (+435T) caused the thermostability to drop to a level that almost matched N1-WA01. Conversely, deleting this codon (−435T) in N1-WA01 increased the thermostability to the level of N1-NY09 (Fig. 6E). However, the analysis of the other mutants indicated that the stability changes are more associated with the insertion (*see* +436A) for N1-NY09 and deletion (*see* −437I) for N1-WA01 than the glycan addition or removal (Fig. 6E). This implies that structural changes imparted by the codon insertion or deletion may affect N1 stability by altering the oligomeric assembly that creates the central Ca^2+^ binding site. We then tested this more broadly by examining the conserved head glycan site mutants. Interestingly, all N1-MI15 mutants lacking the N-X-T site at Asn146 (148A) possess decreased thermostability, indicating that both N-X-T head glycosylation sites contribute to N1 thermostability (Fig. 6F).

### Analysis of N-linked glycan sites encoded by the NA from H3N2 IAVs

H3N2 IAVs commonly circulate together with H1N1 IAVs in the human population. Therefore, we also analysed the subtype 2 (N2) sequences from H3N2 IAVs. In contrast to N1, the avian, swine and human N2 sequences all vary in the number of predicted *N*-linked glycosylation sites, with swine N2s having the most, followed by the human and avian N2s (Fig. 7A, left panel). Surprisingly, almost all N2 sequences were found to contain two sites in the stalk (Fig. 6A, middle panel), resulting in the head domain being responsible for the variation in the number of N2 glycosylation sites (Fig. 7A, right panel).

**Figure 7.**
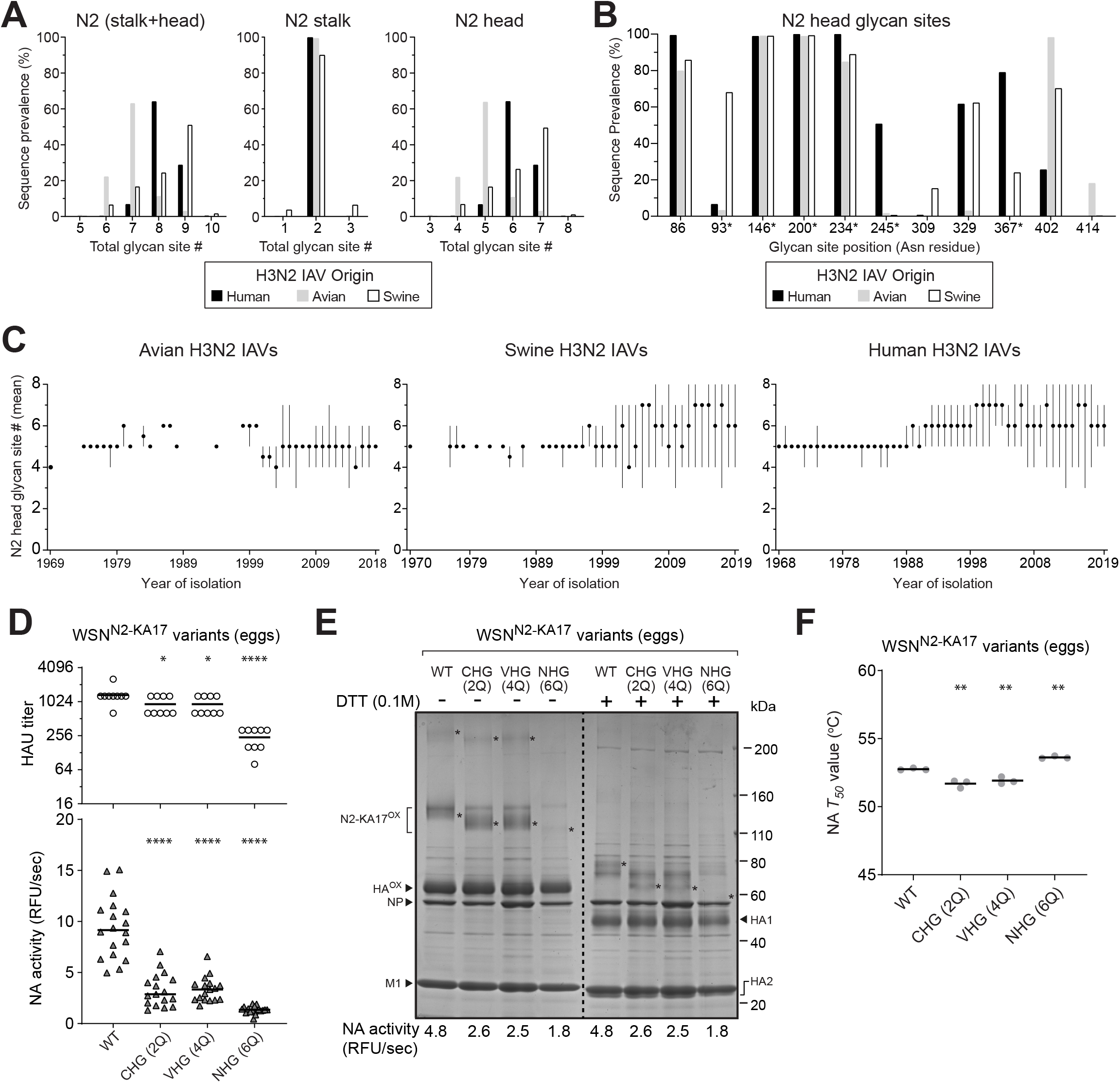
Variation in the NA glycan sites from H3N2 IAVs by species of origin and time. **A.** Graphs showing the prevalence of the NA sequences from human (n=24184), swine (n=3484), and avian (n=411) H3N2 IAVs that possess the indicated number of glycosylation sites in the N2 stalk and head domain (**left panel**), the stalk alone (**middle panel**) and the head domain alone (**right panel**). **B.** The prevalence of the most frequent glycosylation sites in the N2 head domain sequences is shown based on the H3N2 IAV strain species origin. The glycan site position refers to the Asn (N) in the N-X-S/T sequence. Sites with an N-X-T sequence have an asterisk. **C.** Graphs displaying the mean number of glycosylation sites in the N2 head domain with respect to the year the avian (**left panel**), swine (**middle panel**), and human (**right panel**) H3N2 IAVs were isolated. Filled circles correspond to the mean and lines show the range in the site number for the sequence sets from each year. Sequences were obtained from the NCBI Influenza Database. **D.** Scatter plot of the HAU titers and NA activities from the allantoic fluid of individual eggs infected with recombinant WSN viruses carrying N2-KA17 (WT) or mutants that contain only the conserved head glycan sites (CHG 2Q), variable head glycan sites (VHG 4Q), or no head glycan sites (NHG 6Q). Data from the second passage is shown with the median. *P* values (95% CI) were determined with respect to WT by a one-way ANOVA. **E.** The indicated recombinant WSN^N2-KA17^ viruses were isolated from allantoic fluid by centrifugation, separated by SDS-PAGE, and visualized by Coomassie staining. Prior to loading ∼5μg of total protein was treated or untreated with DTT. The NA activity for each virus was measured using equal protein amounts. Asterisks mark bands corresponding to N2-KA17 dimers and tetramers (−DTT) or reduced monomers (+DTT). **F.** NA *T*_50_ temperatures were determined for the indicated WSN reassortant viruses in PBS pH 7.2 with 1mM CaCl_2_. NA activity at 37°C set to 100%. The line is the mean and the *P* values (95% CI) were calculated with respect to WT by a one-way ANOVA.

Positional analysis showed that four of the *N*-linked glycosylation sites are highly conserved in the N2s and that many variable sites showed a bias based on the species of origin of the strain (Fig. 7B). Of the four conserved sites, three (Asn86, Asn146 and Asn234) are nearly identical to N1 and the fourth (Asn200) is located on the side near the dimer interface. In contrast to N1, N-X-T is the most prevalent glycosylation sequence in the N2 head as it is used for three of the conserved sites and three of the common variable sites (Fig. 7B). The temporal analysis showed that most avian N2s carry five sites, whereas the swine and human N2 head domains increased from five to six and seven sites in the more recent H3N2 isolates (Fig. 7C). These observations suggest that N2s may require more head domain glycans for folding; can accommodate more glycans on the head domain; and/or that the N2 head domain is under more selection pressure than N1.

### Contributions of the N2 head glycan sites to viral replication, incorporation and stability

To examine the functional contributions of the N2 head glycosylation sites, several mutants were created using a NA (N2-KA17) from a recently recommended H3N2 vaccine strain (A/Kansas/14/2017) and rescued with a WSN backbone. Following passaging in eggs, lower HAU titres and NA activity levels were obtained for the viruses containing the N2-KA17 mutants with either the four conserved head glycosylation sites (CHG 2Q) or the two variable glycosylation sites (VHG 4Q) at Asn245 and Asn367, and these values decreased further with the no head glycosylation site (NHG 6Q) mutant (Fig. 7D). Upon analysis of the isolated virions, oxidized N2-KA17 dimers were readily apparent for WT that can be reduced by DTT (Fig. 7E). Similar faster migrating bands were observed for the CHG 2Q and VHG 4Q mutants, but the band corresponding to the NHG 6Q mutant was faint. In line with these results, the viruses carrying the CHG 2Q and VHG 4Q mutants possessed ∼50% of the NA activity levels found in the virus containing N2-KA17 WT, whereas the NHG 6Q mutant virus possessed ∼35% (Fig. 7E). Based on the relatively high retention of N2 when the head glycan sites were absent, we examined the stability of the N2 mutants. In contrast to N1, the thermostability of N2 did not significantly change upon removal of the head glycans (Fig. 7F), indicating that glycan addition and removal has a more minimal structural impact on N2.

## DISCUSSION

In this study, sequence-based analyses were combined with several experimental approaches to examine the potential functions of the *N*-linked glycans on NA from H1N1 and H3N2 IAVs. Our results show that three glycosylation sites (Asn88, Asn146 and Asn235) are well-conserved on the N1 head domain and that N2 possesses four conserved sites (Asn86, Asn146, Asn200 and Asn234), which are similar in position. Based on the available sequences, it appears that variable glycosylation sites on the NA head domain are primarily found on human H1N1 IAV strains and that nucleotide substitutions, insertions and/or deletions are likely responsible for the temporal nature of these sites, together with reassortant events involving the NA gene segment [39–41]. In contrast, the variable sites on the NA head domain in H3N2 strains are not exclusive to the species of origin and these mainly appear to result from nucleotide substitutions. Despite the position conservation, none of the head domain glycosylation sites were essential for viral replication in MDCK cells, or eggs, indicating IAVs would not be significantly impacted by inefficient recognition of these sites. In line with a potential role in NA maturation [21], viral growth defects were observed when all the conserved sites on the NA head domain were absent and these coincided with a decrease in the virion incorporation of NA. However, NA activity was detected in all the viruses containing mutations in the head glycosylation sites, indicating a portion of NA can properly mature when one or more of the conserved head domain sites are absent, raising the question of why these sites are conserved in nature.

Although the results primarily focused on H1N1 and H3N2 IAVs, we also found that the conserved glycosylation sites on the N1 head domain (Asn88, Asn146 and Asn235) are also highly prevalent in avian and swine IAV strains carrying a HxN1 subtype combination, where x is any of the other sixteen HA subtypes. In addition, the conserved sites in the N2 head domain (Asn86, Asn146, Asn200 and Asn234) also exist at a high frequency in human H2N2 strains, swine HxN2 and avian HxN2 strains, but in the latter two, lower conservation was observed for the Asn86 and Asn234 sites. However, only the Asn146 site is conserved in all other avian NA subtypes (N3-N9), and the highly prevalent sites in the head domains of these subtypes vary in number from two to five and in position, indicating the function of NA head glycans may differ between subtypes.

Inefficient recognition of the glycosylation sites may be one reason why multiple sites are conserved, as growth defects were only clear when multiple conserved glycan sites were absent. Along these lines, some H1N1 strains that lack one of the three conserved head glycosylation sites have been found, and only the Asn146 site is the more efficiently recognized sequence N-X-T in H1N1 IAVs and is also present in all other NA subtypes at this position [43]. However, three of the conserved N2 head domain sites contain the N-X-T sequence (Asn146, Asn200 and Asn234) and the gel shifts that were observed indicate the different sites are generally recognized. These observations suggest that the individual sites could provide a subtle increase in the replication efficiency of H1N1 IAVs that was not detected by our assays, but it is equally plausible that the conserved sites provide a growth or immunogenicity advantage *in vivo*, as has been reported for several other viral glycoproteins [45–47].

In support of a possible role *in vivo*, the two N-X-T sites on the N1 head domain (Asn146 and Asn434) both affected the enzymatic properties and the conserved site at Asn146 has previously been reported to possess a unique, wide array, of branched glycan structures [35]. The sites are also proximal to one another and near the central Ca^2+^ binding site on top of the N1 tetramer, indicating the glycan or the site may alter the conformation dependent affinity for this Ca^2+^, which is a major NA stability determinant [26]. In line with this interpretation, less significant stability affects were observed when the variable N-X-T site was inserted into a N1 (NA/CA09) with a lower central Ca^2+^ binding site affinity (data not shown). Currently, we cannot investigate the structural consequence of the insertion more directly because no structure is available for a human N1 head domain between 1948-2009, which contains the amino acid insertion resulting in the additional glycosylation site.

An interesting observation is that N1 sequences predominantly utilize the N-X-S consensus glycosylation site, whereas N2 sequences are significantly biased towards N-X-T sites and some sites show species specific X residues. This suggests that recognition is more critical for N2 than N1, and that the conserved N1 sites are required for a specific function such as limiting epitope access. However, an enzyme-linked lectin (ELLA) analysis [48] of viruses containing N1-NY09 with and without the insertion and Asn434 glycan addition showed no difference in reactivity to a ferret antiserum raised against an NA (N1-CA09) almost identical to N1-NY09. Conversely, N1-WA01 did not gain reactivity against the same antiserum when the deletion was introduced, indicating that the antigenic changes between these two strains is not related to the removal or addition of the Asn434 glycosylation site in the human N1 head domain (data not shown). While this was somewhat t surprising, the region surrounding the central Ca^2+^-binding site on the N1 tetramer has previously been shown to be highly conserved [26], indicating that it may not be subject to significant selection pressure or that antibodies binding to this region do not negatively impact viral replication.

Assigning functions to *N*-linked glycans on viral glycoproteins is difficult, as a role in maturation does not exclude an additional role in altering surface epitopes. The observation that the number of glycan sites on the N1 head domain have both increased and decreased over time, whereas those on N2 have primarily increased in number, suggests that glycan site addition may have subtype-dependent effects. Supporting this possibility, N2 has a higher prevalence of head glycosylation sites and variable sites, and the N1 variable site at Asn386, which was introduced during the 2009 pandemic, was quickly lost in the circulating strains. There also are additional glycosylation sites with a low frequency in the database that we did not examine, indicating some may be advantageous in specific populations. In this study, we demonstrated that the glycosylation sites on the NA head domain are required for efficient virion incorporation and replication, indicating mutations in these sites are likely useful for creating attenuated IAV strains. These mutant strains can also be used for future studies examining how the conserved N-linked glycan sites on NA contribute to IAV viability, antigenicity, antibody binding and transmissibility in addition to their likely function during maturation.

## MATERIALS AND METHODS

### Reagents and antibodies

Dulbecco’s Modified Eagles Medium (DMEM), fetal bovine serum (FBS), L-glutamine, penicillin/streptomycin (P/S), Opti-MEM (OMEM), anti-goat IgG HRP-linked secondary antibody, Simple Blue Stain, Novex 4-12% Tris-Glycine SDS-PAGE gels, lipofectamine and dithiothreitol (DTT) were all purchased from Thermo Fisher Scientific. 2’-(4-methylumbelliferyl)-α-d-*N*-acetylneuraminic acid (MUNANA) was obtained from Cayman Chemical. Anti-rabbit IgG HRP-linked secondary antibody and 0.45-μm polyvinylidene difluoride (PVDF) membrane were obtained from GE healthcare. Specific-Pathogen-Free (SPF) eggs and turkey red blood cells (TRBCs) were purchased from Charles River Labs and the Poultry Diagnostic and Research Center (Athens, GA), respectively. Rabbit antisera against NA was generated by Agrisera (Sweden) using NA-WSN33 residues 35–453 isolated from *E. coli* inclusion bodies. Polyclonal goat antisera against influenza virus HAs from A/California/04/2009 (NR-15696) and A/Fort Monmout/1/1947 (NR-3117) were both obtained from BEI Resources, NIAID, NIH.

### Plasmids and constructs

The eight reverse genetics (RG) plasmids encoding the PR8 and WSN33 gene segments were provided by Dr. Robert Webster (St. Jude Children’s Research Hospital). The RG plasmid containing NA (N1-NY09) from the H1N1 strain A/New York/18/2009 was described previously [26]. The NA gene segments from the strains A/Waikato/7/2001 (N1-WA01), A/Michigan/45/2015 (N1-MI15), A/Kansas/14/2017 (N2-KA17) and the CHG 2Q, VHG 4Q, and NHG 6Q were all synthesized (Eurofins Genomics or GeneScript) and used to generate the RG plasmids as follows. The pHW2000 plasmid back bone [49] and the NA gene segments were amplified by PCR using forward and reverse primers (Table 1) with complementary NA 5’ and 3’ UTR overhangs that direct the recombination upon transformation into *E. coli* [50]. The N1-BR18 RG plasmid was generated by cloning the NA gene segment from the H1N1 strain A/Brisbane/02/2018 IVR-190 grown in SPF eggs into the plasmid pHW2000 following PCR amplification [51]. Mutations in the N1-MI15 head domain (S90A, S90P, T148A and lineN235K) and the N1-BR18 head domain (N86Q, N146Q and N235), codon insertions in N1-NY09 (+435T and + 436A), codon deletions in N1-WA01 (−435T and −437I), and the substitutions in N1-NY09 (I436T) and N1-WA01 (T436A) were all made with site-directed mutagenesis primers (Table 2) using the respective NA RG plasmid as a template. All constructs were confirmed by sequencing (Eurofins MWG Operon or the FDA core facility).

**Table 1.**
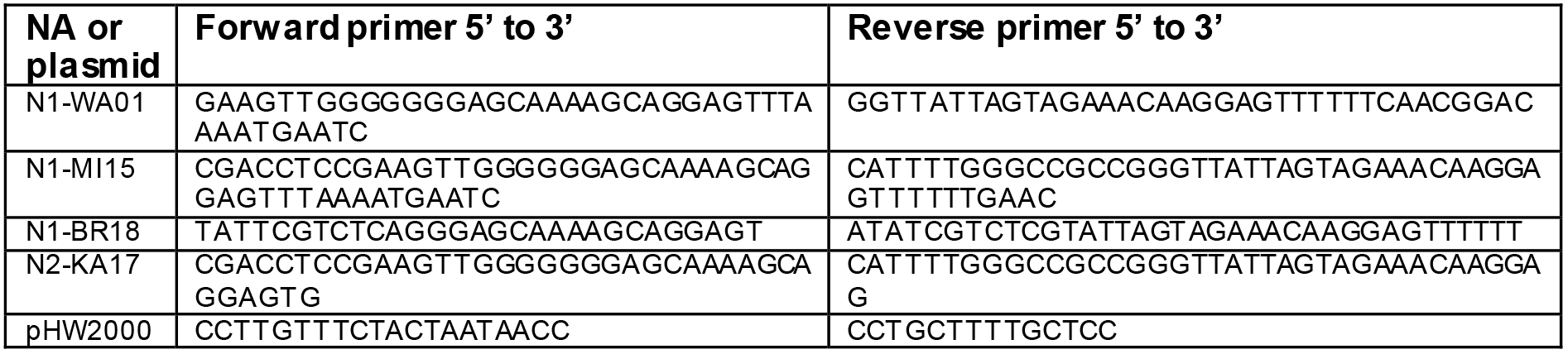
Primers used for inserting the NA gene segments into the pHW2000 plasmid.

**Table 2.**
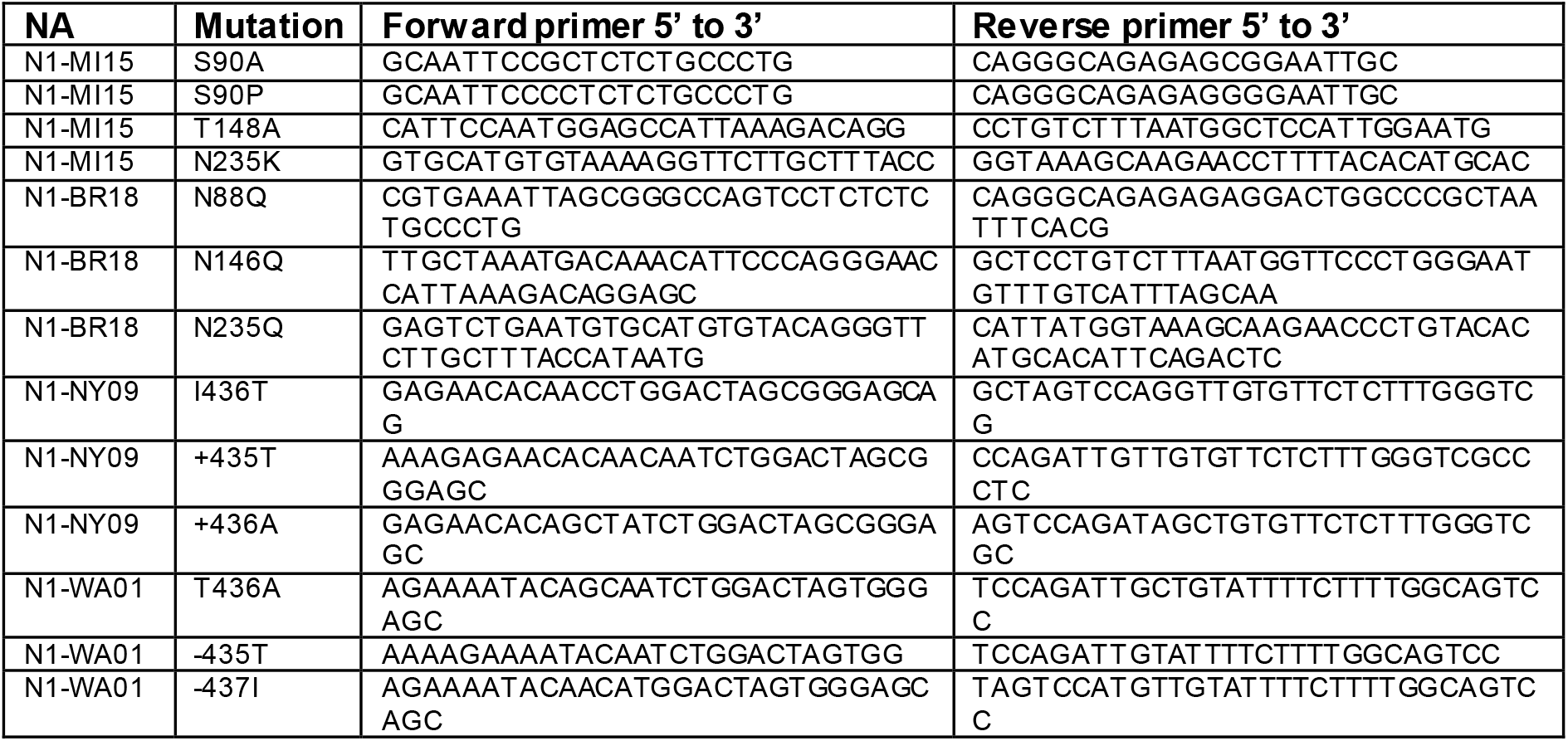
Primers used for introducing the site-directed mutations in the NA head domain.

### Cell culture and viral reverse genetics

Madin-Darby canine kidney 2 (MDCK.2; CRL-2936) cells and HEK 293T/17 cells (CRL-11268), obtained from LGC Standards, were cultured in DMEM containing 10% FBS and 100 U/ml of P/S in a 37 °C atmosphere with 5% CO_2_ and ∼95% humidity. Reassortant viruses carrying N1-MI15, N1-NY09 or N1WA01 variants were generated by 8-plasmid reverse genetics using the indicated NA and the complimentary seven ‘backbone’ gene segments of WSN33 as previously described [26]. Reassortant viruses carrying N1-BR18 or N2-KA17 variants were generated by 8-plasmid reverse genetics using the seven ‘backbone’ gene segments of WSN33 or PR8 as follows using 6-well plates. One day prior, 1.2×10^6^ 293T cells and 1.2×10^6^ MDCK.2 cells were plated per well using 3 ml DMEM with 10% FBS. The next day, the medium was replaced with 2 ml of OMEM, the eight plasmids were added to 200 μl of OMEM at a concentration of 1 μg per plasmid, combined with 18 μl of lipofectamine and the mixture was incubated 45 min at room temperature. The cell medium was removed, the mixture was added to one well and the dish was incubated 5 min at 37 °C before 800 μl OMEM was added to each well. Approximately 24 h post-transfection 1ml OMEM containing 4 μg/ml TPCK trypsin was added to each well. Culture medium was harvested between 72-96 h post-transfection, clarified by centrifugation (2000 × g; 5 min) and passaged using SPF eggs or MDCK.2 cells.

### Viral Passaging in MDCK.2 cells and SPF eggs

For cell passaging, one day after seeding 1 × 10^6^ MDCK.2 cells on a 6 cm dish, the culture medium was removed, and the cells were washed with 1 ml infection medium (IM) comprised of DMEM, 0.3% BSA, 0.1% FBS, and P/S. Each dish then received 2 ml of cold IM containing 10 μl of the clarified viral reverse genetics medium and was rocked at 4 °C for 30 min. The inoculation medium was then removed, cells were washed with 1 ml of IM, 5 ml of IM containing 10 μg/ml TPCK-trypsin was added and the dish was placed at 37 °C. The culture medium was harvested at the peak of cytopathic effects (∼48-72 h post-infection) and clarified by centrifugation (2000 × g; 5 min) prior to analysis, storage, or sedimentation. Passaging in SPF eggs was carried out by inoculating 100 μl of clarified viral reverse genetics medium into 9-11 day old embryonated eggs and incubating them for 3 days at 33 °C. Following the incubation, eggs were chilled at 4 °C for 2 h and the allantoic fluid from each egg was harvested, clarified by centrifugation (2000 × g; 5 min) and stored in aliquots at −80 °C. Viruses in the allantoic fluid from the first passage were then diluted (1:1000) in sterile PBS and 100 μl was used to inoculate the 9-11 day old embryonic eggs. For each virus, groups of eight or seven eggs were used, the allantoic fluid from each egg was harvested individually and clarified by centrifugation (2000 × g; 5 min) prior to analysis or sedimentation.

### Viral sedimentation and sucrose gradient isolation

Clarified virus-containing culture medium (∼8 ml) or allantoic fluid (∼28 ml) were added to ultracentrifuge tubes and a sucrose cushion (25% w/v sucrose, PBS pH 7.2 and 1 mM CaCl_2_) equal to ∼15% of the sample volume was layered under each sample. Virions were then isolated by sedimentation (100,000 × g; 45 min) at 4 °C and the supernatant was aspirated. Cell produced viral pellets were resuspended in 200 μl PBS pH 7.4 and 1 mM CaCl_2_ and NA activity was analysed prior to immunoblotting. Egg produced virions were resuspended in 200 μl of PBS pH 7.2 containing 1 mM CaCl_2_. For sucrose gradient isolations, the resuspension solution containing 12.5% w/v sucrose was layered on top of a discontinuous gradient containing four 8.5 ml sucrose layers (60% w/v, 45% w/v, 30% w/v and 15% w/v sucrose in PBS pH 7.2 and 1 mM CaCl_2_) and centrifuged at 100,000 × g for 2 h at 4 °C. Fractions were isolated from top to bottom, the density was determined with a refractometer and those corresponding to 30-50% w/v sucrose were pooled, mixed with 2 volumes of PBS pH 7.2 and 1 mM CaCl_2_, and the virions were sedimented (100,000 × g; 45 min). The supernatant was discarded, and the viral pellet was resuspended in 250 μl PBS pH 7.2 containing 1 mM CaCl_2_. Total protein concentrations in resuspended viral pellets were all determined with a BCA protein assay kit (Pierce) using the 96-well plate protocol. The average value was determined from the 1:2 and 1:4 sample dilutions and each sample was adjusted to a final concentration of 1 mg/ml using PBS pH 7.2 containing 1 mM CaCl_2_.

### HAU titre, NA activity and thermostability measurements

HAU titres were determined using a 96-well plate and 0.5% TRBCs in PBS pH 7.2. Briefly, 90 μl of PBS pH 7.2 was added to the first column and 50 μl to remaining columns. From each infected egg, 10 μl of allantoic fluid was added to the first column creating a 1:10 dilution. A two-fold serial dilution was made by transferring 50 μl from each column to the subsequent column and 50 μl of 0.5% TRBCs were added to each well. The plate was incubated 30 min at room temperature and the HAU titre was determined as the last well where agglutination was observed. For sialidase activity measurements, equal amounts of clarified virus-containing medium, allantoic fluid or sedimented viral samples were brought up to 195 μl in reaction buffer (0.1 M KH_2_PO_4_ pH 6.0 and 1 mM CaCl_2_), transferred to a 96-well black clear bottom plate (Corning) and incubated at 37 °C for 15 min. Reactions were then initiated by adding 5 μl of 2 mM MUNANA and the fluorescence was measured with either a SpectraMax Gemini EM plate reader or a Cytation 5 (Biotek) using 365 nm excitation and 450 nm emission wavelengths. NA thermostability was determined by exposing equal amounts of clarified virus-containing media to temperatures ranging from 37 °C to 64 °C for 10 min and measuring the residual sialidase activity as previously described [26].

### SDS-PAGE, Coomassie staining and immunoblotting

Sedimented viral samples containing equal resuspension volumes or the indicated total protein amounts were mixed with Laemmli sample buffer that contained 0.1M DTT as indicated. Samples were then heated 37 °C or 50 °C for 10 min and resolved by either 7.5 % (α-NA), 11 % (α-HA) or 4-12 % (α-NA, α-HA and Coomassie) Tris-Glycine SDS-PAGE gels. Gels were Coomassie stained using simple blue or transferred to a 0.45-μm pore PVDF membrane at 15 V for 1 h. PVDF membranes were blocked with milk/PBST (3% nonfat dry milk, PBS, pH 7.4, 0.1% Tween 20) for 30 min and processed using standard immunoblotting protocols with the indicated antibodies and the appropriate HRP-linked secondary antibody. Immunoblots were developed with the SuperSignal West Femto kit (ThermoFischer) and imaged using an Azure C600 or a Syngene G Box.

### Analysis of NA glycosylation sites and in silico glycosylation models

Complete NA protein sequences from H1N1 and H3N2 IAVs of human, avian and swine origin were downloaded from The Influenza Virus Resource at the National Center for Biotechnology Information [52]. Each group was aligned using MAFFT v7.311 with default progressive method (FFT-NS-2). Mislabeled sequences were manually removed. The final NA data sets consist of 18966 sequences from human H1N1 strains (1918-2019/11/20), 630 sequences from avian H1N1 strains (1976-2018/11/16), 4949 sequences from swine H1N1 strains (1930-2019/12/20), 24184 sequences from human H3N2 strains (1968-2019/11/09), 411 sequences from avian H3N2 strains (1969-2018/10/13), and 3484 sequences from swine H3N2 strains (1970-2019/12/19). Potential glycosylation sites (N-X-S/T-X), where X represents every amino acid except for Pro, were located using a Python script. N1 2009 pandemic-like amino acid numbering was used for both N1 and N2, the head domain was set to begin at amino acid residue 82 and the stalk was designated as amino acid residues 35 to Tetrameric N1 models were created from the A/Michigan/45/2015 primary sequence using SWISS-MODEL (https://swissmodel.expasy.org), based on an available 2009 pandemic-like N1 head domain structure (PDBID: 5NWE) [53], and the glycans were added *in silico* using Glyprot (http://www.glycosciences.de/modeling/glyprot/php/main.php). The modeled glycan structures were chosen based on previous work [35] with glycan number 9141 (glcp) being used for Asn386 and 8714 (2 glcnac) for all other sites.

## ACKNOWLEDGEMENTS

We would like to thank Tahir Malik (FDA), Hongquan Wan (FDA) and Daniel Hebert (University of Massachusetts-Amherst) for critically reading the manuscript and offering several helpful suggestions. This work was supported by in part by grants from the Swedish Research Council K2015-57-21980-04-4 and the Carl Trygger Foundation CTS17:111, as well as federal funds from the NIAID, National Institutes of Health, Department of Health and Human Services, under CEIRS contract number HHSN272201400005C.

## REFERENCES

1. Mochizuki, K., et al., Two N-linked glycans are required to maintain the transport activity of the bile salt export pump (ABCB11) in MDCK II cells. Am J Physiol Gastrointest Liver Physiol, 2007. 292(3): p. G818–28.

2. Hanson, S.R., et al., The core trisaccharide of an N-linked glycoprotein intrinsically accelerates folding and enhances stability. Proc Natl Acad Sci U S A, 2009. 106(9): p. 3131–6.

3. Tokhtaeva, E., et al., N-glycan-dependent quality control of the Na,K-ATPase beta(2) subunit. Biochemistry, 2010. 49(14): p. 3116–28.

4. Hebert, D.N., et al., The intrinsic and extrinsic effects of N-linked glycans on glycoproteostasis. Nat Chem Biol, 2014. 10(11): p. 902–10.

5. Cai, X., et al., The importance of N-glycosylation on beta3 integrin ligand binding and conformational regulation. Sci Rep, 2017. 7(1): p. 4656.

6. Ohtsubo, K., et al., Dietary and genetic control of glucose transporter 2 glycosylation promotes insulin secretion in suppressing diabetes. Cell, 2005. 123(7): p. 1307–21.

7. Wang, X., et al., Core fucosylation regulates epidermal growth factor receptor-mediated intracellular signaling. J Biol Chem, 2006. 281(5): p. 2572–7.

8. Goetze, A.M., et al., High-mannose glycans on the Fc region of therapeutic IgG antibodies increase serum clearance in humans. Glycobiology, 2011. 21(7): p. 949–59.

9. Liu, L., Antibody glycosylation and its impact on the pharmacokinetics and pharmacodynamics of monoclonal antibodies and Fc-fusion proteins. J Pharm Sci, 2015. 104(6): p. 1866–1884.

10. Hebert, D.N., B. Foellmer, and A. Helenius, Glucose trimming and reglucosylation determine glycoprotein association with calnexin in the endoplasmic reticulum. Cell, 1995. 81(3): p. 425–33.

11. Shi, X. and R.M. Elliott, Analysis of N-linked glycosylation of hantaan virus glycoproteins and the role of oligosaccharide side chains in protein folding and intracellular trafficking. J Virol, 2004. 78(10): p. 5414–22.

12. Braakman, I. and E. van Anken, Folding of viral envelope glycoproteins in the endoplasmic reticulum. Traffic, 2000. 1(7): p. 533–9.

13. Skehel, J.J., et al., A carbohydrate side chain on hemagglutinins of Hong Kong influenza viruses inhibits recognition by a monoclonal antibody. Proc Natl Acad Sci U S A, 1984. 81(6): p. 1779–83.

14. Wei, X., et al., Antibody neutralization and escape by HIV-1. Nature, 2003. 422(6929): p. 307–12.

15. Lennemann, N.J., et al., Comprehensive functional analysis of N-linked glycans on Ebola virus GP1. mBio, 2014. 5(1): p. e00862–13.

16. Walls, A.C., et al., Glycan shield and epitope masking of a coronavirus spike protein observed by cryo-electron microscopy. Nat Struct Mol Biol, 2016. 23(10): p. 899–905.

17. Wan, H., et al., The neuraminidase of A(H3N2) influenza viruses circulating since 2016 is antigenically distinct from the A/Hong Kong/4801/2014 vaccine strain. Nat Microbiol, 2019. 4(12): p. 2216–2225.

18. Hebert, D.N., et al., The number and location of glycans on influenza hemagglutinin determine folding and association with calnexin and calreticulin. Journal of Cell Biology, 1997. 139(3): p. 613–623.

19. Daniels, R., et al., N-linked glycans direct the cotranslational folding pathway of influenza hemagglutinin. Mol Cell, 2003. 11(1): p. 79–90.

20. Molinari, M., et al., Contrasting functions of calreticulin and calnexin in glycoprotein folding and ER quality control. Mol Cell, 2004. 13(1): p. 125–35.

21. Wang, N., et al., The cotranslational maturation program for the type II membrane glycoprotein influenza neuraminidase. J Biol Chem, 2008. 283(49): p. 33826–37.

22. Gamblin, S.J. and J.J. Skehel, Influenza hemagglutinin and neuraminidase membrane glycoproteins. J Biol Chem, 2010. 285(37): p. 28403–9.

23. Yoon, S.W., R.J. Webby, and R.G. Webster, Evolution and ecology of influenza A viruses. Curr Top Microbiol Immunol, 2014. 385: p. 359–75.

24. Morens, D.M., J.K. Taubenberger, and A.S. Fauci, The persistent legacy of the 1918 influenza virus. N Engl J Med, 2009. 361(3): p. 225–9.

25. Rajao, D.S., A.L. Vincent, and D.R. Perez, Adaptation of Human Influenza Viruses to Swine. Front Vet Sci, 2018. 5: p. 347.

26. Wang, H., et al., Structural restrictions for influenza neuraminidase activity promote adaptation and diversification. Nature Microbiology, 2019.

27. Bos, T.J., A.R. Davis, and D.P. Nayak, NH2-terminal hydrophobic region of influenza virus neuraminidase provides the signal function in translocation. Proc Natl Acad Sci U S A, 1984. 81(8): p. 2327–31.

28. Paterson, R.G. and R.A. Lamb, Conversion of a class II integral membrane protein into a soluble and efficiently secreted protein: multiple intracellular and extracellular oligomeric and conformational forms. J Cell Biol, 1990. 110(4): p. 999–1011.

29. Bucher, D.J. and E.D. Kilbourne, A 2 (N2) neuraminidase of the X-7 influenza virus recombinant: determination of molecular size and subunit composition of the active unit. J Virol, 1972. 10(1): p. 60–6.

30. Varghese, J.N., W.G. Laver, and P.M. Colman, Structure of the influenza virus glycoprotein antigen neuraminidase at 2.9 A resolution. Nature, 1983. 303(5912): p. 35–40.

31. Dou, D., et al., Influenza A Virus Cell Entry, Replication, Virion Assembly and Movement. Frontiers in Immunology, 2018. 9(1581).

32. Colman, P.M., J.N. Varghese, and W.G. Laver, Structure of the catalytic and antigenic sites in influenza virus neuraminidase. Nature, 1983. 303(5912): p. 41–4.

33. Nordholm, J., et al., Polar residues and their positional context dictate the transmembrane domain interactions of influenza a neuraminidases. J Biol Chem, 2013. 288(15): p. 10652–60.

34. Dou, D., et al., Type II transmembrane domain hydrophobicity dictates the cotranslational dependence for inversion. Mol Biol Cell, 2014. 25(21): p. 3363–74.

35. She, Y.M., et al., Topological N-glycosylation and site-specific N-glycan sulfation of influenza proteins in the highly expressed H1N1 candidate vaccines. Sci Rep, 2017. 7(1): p. 10232.

36. da Silva, D.V., et al., Assembly of subtype 1 influenza neuraminidase is driven by both the transmembrane and head domains. J Biol Chem, 2013. 288(1): p. 644–53.

37. da Silva, D.V., et al., The influenza virus neuraminidase protein transmembrane and head domains have coevolved. J Virol, 2015. 89(2): p. 1094–104.

38. Saito, T., G. Taylor, and R.G. Webster, Steps in maturation of influenza A virus neuraminidase. J Virol, 1995. 69(8): p. 5011–7.

39. Sun, S., et al., Glycosylation site alteration in the evolution of influenza A (H1N1) viruses. PLoS One, 2011. 6(7): p. e22844.

40. Sun, S., et al., Prediction of biological functions on glycosylation site migrations in human influenza H1N1 viruses. PLoS One, 2012. 7(2): p. e32119.

41. Kim, P., et al., Glycosylation of Hemagglutinin and Neuraminidase of Influenza A Virus as Signature for Ecological Spillover and Adaptation among Influenza Reservoirs. Viruses, 2018. 10(4).

42. Shakin-Eshleman, S.H., S.L. Spitalnik, and L. Kasturi, The amino acid at the X position of an Asn-X-Ser sequon is an important determinant of N-linked core-glycosylation efficiency. J Biol Chem, 1996. 271(11): p. 6363–6.

43. Mellquist, J.L., et al., The amino acid following an asn-X-Ser/Thr sequon is an important determinant of N-linked core glycosylation efficiency. Biochemistry, 1998. 37(19): p. 6833–7.

44. Eichelberger, M.C. and H. Wan, Influenza neuraminidase as a vaccine antigen. Curr Top Microbiol Immunol, 2015. 386: p. 275–99.

45. Lavie, M., X. Hanoulle, and J. Dubuisson, Glycan Shielding and Modulation of Hepatitis C Virus Neutralizing Antibodies. Frontiers in Immunology, 2018. 9(910).

46. Quinones-Kochs, M.I., L. Buonocore, and J.K. Rose, Role of N-linked glycans in a human immunodeficiency virus envelope glycoprotein: effects on protein function and the neutralizing antibody response. J Virol, 2002. 76(9): p. 4199–211.

47. Leemans, A., et al., Removal of the N-Glycosylation Sequon at Position N116 Located in p27 of the Respiratory Syncytial Virus Fusion Protein Elicits Enhanced Antibody Responses after DNA Immunization. Viruses, 2018. 10(8).

48. Gao, J., L. Couzens, and M.C. Eichelberger, Measuring Influenza Neuraminidase Inhibition Antibody Titers by Enzyme-linked Lectin Assay. J Vis Exp, 2016(115).

49. Hoffmann, E., et al., A DNA transfection system for generation of influenza A virus from eight plasmids. Proc Natl Acad Sci U S A, 2000. 97(11): p. 6108–13.

50. Mellroth, P., et al., LytA, major autolysin of Streptococcus pneumoniae, requires access to nascent peptidoglycan. J Biol Chem, 2012. 287(14): p. 11018–29.

51. Sandbulte, M.R., et al., A miniaturized assay for influenza neuraminidase-inhibiting antibodies utilizing reverse genetics-derived antigens. Influenza Other Respir Viruses, 2009. 3(5): p. 233–40.

52. Bao, Y., et al., The influenza virus resource at the National Center for Biotechnology Information. J Virol, 2008. 82(2): p. 596–601.

53. Pokorna, J., et al., Kinetic, Thermodynamic, and Structural Analysis of Drug Resistance Mutations in Neuraminidase from the 2009 Pandemic Influenza Virus. Viruses, 2018. 10(7).

